# SLFL Genes Participate in the Ubiquitination and Degradation of *S*-RNase in Self-Compatible Chinese Peach

**DOI:** 10.1101/160267

**Authors:** Qiuju Chen, Dong Meng, Wei Li, Zhaoyu Gu, Hui Yuan, Xuwei Duan, Qing Yang, Yang Li, Tianzhong Li

## Abstract

The gametophytic self-incompatibility (SI) mediated by S-RNase of Rosaceae, Solanaceae and Plantaginaceae, is controlled by two tightly linked genes located at highly polymorphic S-locus: the S-RNase for pistil specificity and the F-box gene (SFB/SLF) for pollen specificity, respectively. The F-box gene of peach (*Prunus persica*) is *S* haplotype-specific F-box (*SFB*). In this study, we selected 37 representative varieties according to the evolution route of peach and identified their S genotypes. We cloned pollen determinant genes mutant *PperSFB1m, PperSFB2m, PperSFB4m* and normal *PperSFB2*, and style determinant genes *S1-RNase, S2-RNase, S2m-RNase* and *S4-RNase.* Mutant *PperSFBs* were translated terminated prematurely because of fragment insertion. Yeast two-hybrid showed that mutant PperSFBs and normal PperSFB2 interacted with all S-RNases. Normal *PperSFB2* was divided into four parts: box, box-V1, V1-V2 and HVa-HVb. Protein interaction analyses showed that the box portion did not interact with S-RNases, both of the box-V1 and V1-V2 had interactions with S-RNases, while the hypervariable region of *PperSFB2* HVa-HVb only interacted with S2-RNase. Bioinformatics analysis of peach genome revealed that there were other F-box genes located at S-locus, and of which three F-box genes were specifically expressed in pollen, namely *PperSLFL1, PperSLFL2* and *PperSLFL3*, respectively. Phylogenetic analysis showed that PperSFBs and PperSLFLs were classified into two different clades. Yeast two-hybrid analysis revealed that as with PperSFBs, the three F-box proteins interacted with PperSSK1. Yeast two-hybrid and BiFC showed that PperSLFLs interacted with S-RNases with no allelic specificity. In vitro ubiquitination assay showed that PperSLFLs could tag ubiquitin molecules to PperS-RNases. In all, the above results suggest that three *PperSLFLs* are the appropriate candidates for the ‘general inhibitor’, which would inactivate the S-RNases in pollen tubes, and the role of three PperSLFL proteins is redundant, as S-RNase repressors involved in the self-incompatibility of peach.

## Introduction

Self-incompatibility (GSI) is the most widely distributed breeding system that allows the pistil to reject genetically related pollen and promotes out-crossing in flowering plants (de Nettancourt, 2001). Many species in Solanaceae, Rosaceae, and Plantaginaceae exhibit S-RNase-based GSI which is controlled by the complex S-locus which contains at least two genes, one gene is pistil-part, a highly polymorphic *S* gene encoding extracellular ribonuclease called S-RNase, and one is pollen-part specific *S* gene, which is tightly linked to the S-RNase. The tightly linked genetic unit of the pistil S allele (S-RNase) and pollen S allele is called S haplotype. The pollen *S* genes of the S-RNase-based GSI of the above three families are F-box genes called *SLF/SFB* (Entani et al., 2003; Lai et al., 2002; Sijacic et al., 2004; Ushijima et al., 2003; Yamane et al., 2003). The pollen S gene was called *SLF* (S locus F-box) in Solanaceae and Plantaginaceae and *SFB* (S haplotype-specific F-box) in *Prunus* (Sassa et al., 2010; Tao and Iezzoni, 2010; Meng et al., 2011). S-RNase is secreted into style transmitting intercellular space, and non-selectively taken up into cytoplasm of compatible and incompatible pollen tubes elongating in style tissues, and pollen tube elongation is arrested in incompatible crosses, probably because of cytotoxic effects of self S-RNase (Luu et al., 2000; Goldraij et al., 2006; McClure et al., 2011; Boivin et al., 2014). The predominant role of F-box proteins is ubiquitination of proteins, thus possibly SLF/SFB acts as a subunit to generally constitute the SCF complex, an E3 ubiquitin ligase, discriminates between self and non-self S-RNase, and mediate the ubiquitination of non-self S-RNases for degradation by the 26S proteasome (Lechner et al., 2006; Franklin-Tong, 2008; Hua et al., 2008). In this process, how F-box proteins discriminate self and non-self S-RNases in pollen tubes is unknown.

A single S-locus F-box gene is known as *SFB* identified in *Prunus* (Entani et al., 2003; Ushijima et al., 2003; Sonneveld et al., 2005), while multiply *F-box* genes located at the S-locus have been cloned in subtribe Maloideae designated as *F-box brothers* (Sassa et al., 2007; Kubo et al., 2010; Kakui et al., 2011). In subfamily Maloideae (e.g., apple and pear) of Rosaceae, polyploidization breaks SI in pollen but does not affect the pistil (de Nettancourt, 2001). The pistil of ‘Fertility’ (2x) could accept pollen from autotetraploid (4x), but ‘Fertility’ (2x) pollen was rejected by the pistil of autotetraploid (4x) (Crane and Lewis, 1942). Genetic analyses reveal that the breakdown of SI can be explained by ‘competition’ between different S alleles in pollen. But in *Prunus* (subfamily Prunoideae), tetraploidy is not always associated with SC. Sour cherry (*Prunus cerasus*) is a naturally occurring tetraploid species and includes both SI and SC plants (Lansari and Iezzoni, 1990). Genetic analysis of SI sour cherry suggested that heteroallelic diploid pollen tubes are rejected by pistils with cognate S haplotypes (Hauck et al., 2006). Hauck et al. (2006) proposed that the breakdown of SI in tetraploid sour cherry is caused by the accumulation of non-functional S haplotypes and not by competitive interaction in heteroallelic pollen. In Japanese pear, S^4sm^ pollen lacking SFBB1-S^4^ are rejected by compatible S^1^ pistils but accepted by S^3^ and S^5^ pistils (Okada et al., 2004; Okada et al., 2008). On the other hand, loss-of-function of SFBB1-S^5^ had no effect on SI phenotype. Genetic analysis reveals that S^5^ pollen is normally accepted by S^1^, S^3^ and S^4^ pistils (Sassa et al., 2011). The previous fruit set analyses shows that S^5^ pollen is normally compatible with S^2^ and S^9^ pistils and incompatible with S^5^ pistils (Kajiura et al., 1967; 1969; 1974). On the contrary, in *Prunus*, a truncated SFB protein or lacked the SFB gene can confer pollen-part self-compatibility (SC) (Ushijima et al., 2004; Sonneveld et al., 2005; Hauck et al., 2006; Tsukamoto et al., 2006). These findings suggest that S-RNase-based GSI seems to consist of two types in which the mode of action of pollen S is different, a ‘self recognition by a single factor’ system and a ‘non-self recognition by multiple factors’ system (Kakui et al., 2011), and the S-RNase-based GSI of *Prunus* represents ‘self recognition by a single factor’, in which the cytotoxic effect of non-self S-RNases in pollen tubes is inactivated by a ‘general inhibitor’ while the self S-RNase is specifically protected by a ‘blocker’ molecule and degrades RNA of self-pollen to arrest tube growth (Luu et al., 2001; Sonneveld et al., 2005). Although the ‘general inhibitor’ is a hypothetical protein and had been considered to be SLFLs in *Prunus avium* (Matsumoto and Tao, 2016), and in peach, whether the ‘general inhibitor’ is SLFLs as with in *Prunus avium* needs to be studied.

Skp1, Cullin1 (Cul1), Rbx1 and F-box proteins together constitute the SCF complex, E3 ubiquitin ligases. The E3 ubiquitin ligase can make substrate proteins polyubiquitination to degrade by the 26s proteasome system. In the SCF complex, the F-box protein determines substrate specifically, Skp1 serves as an adaptor to connect the variable F-box protein and Cul1 protein, Cul1 forms a core catalytic scaffold with Rbx1, and Rbx1 can bind to E2 and catalyzes the transfer of ubiquitin chains from E2 to the substrate protein to make ubiquitination of substrate proteins (Wu et al., 2000; Zheng et al., 2002; Deshaies and Joazeiro, 2009). SLF/SFBB is shown to be a compoment of the SCF complex to detoxified non-self S-RNases. In petunia, SLF was showed to form the SCF complex with Skp1-like and Cul1-p in pollen(Zhao et al., 2010; Entani et al., 2014; Liu et al., 2014), and in Maloideae, SFBB was also shown to form SCF complex which targeted selectively S-RNase and polyubiquitinated it in vitro (Yuan et al., 2014).

In our study, we selected 37 representative species according to the evolution route of peach, and identified their S genotypes. Through yeast two-hybrid and BiFC analysis, we found that PperSFB2 distinguished self S2-RNase from non-self S-RNases by the C-terminal hypervariable region. According to the genome wide analysis, we cloned three *SLFL* genes in the S-locus on chromosome 6, and did some experiments and analysis to determine whether the function of the *SLFL* genes is the same as that of *Prunus avium.* Our results showed that PperSLFL proteins were in different clade compared with PperSFB proteins, and could participate in self-incompatibility of peach as a subunit of SCF complex.

## Materials and Methods

### Plant Material

37 peach varieties (Supplemental Table 1) were selected from the Zhengzhou Fruit Research Institute, Chinese Academy of Agricultural Sciences, Henan Province, China. Peach organs/tissue samples (leaves, styles and pollen) were collected, frozen in liquid nitrogen and stored at -80°C Ultra-low temperature refrigerator for later use.

### DNA and RNA Extraction

Peach genomic DNA was isolated from young leaves using the CTAB method (Li et al., 2009), and incubated with RNase I (Invitrogen, CA, USA) at 37°C for 2 hours to remove RNA. Total RNA samples were isolated from leaves, styles and pollen using a modified CTAB method (Li et al., 2009) and treated with DNase I (Invitrogen, CA, USA) to remove DNA contamination. RNA was used as template to synthesize first-strand cDNA using the SuperScript reverse transcriptase (Invitrogen, CA, USA) and Oligo-dT primers (According to manufacturer’s instructions, Invitrogen, CA, USA).

### PCR for S Genotype Analysis

Peach genomic DNA was used as templates for PCR with the primers listed in Supplemental Table 2. The primers were designed according to the length of the second intron of the S-RNases. The different S-RNases and S genotypes of peach varieties could be distinguished depending on the size of the amplified fragments.

### Cloning of *PperS-RNases, PperSFBs, PperSLFLs, PperSSK1, PperCUL1* and *PperRbx1*

Pollen cDNA was used as template to clone, *PperSFBs, PperSLFLs, PperSSK1, PperCUL1* and *PperRbx1* and style cDNA was used as template to clone *PperS-RNases* with the gene specific primers listed in Supplemental Table S2. The PCR products were purified and individually ligated to the pMD19T-simple vector (TaKaRa).The constructed vectors were transformed into *E. coli* competent DH5α cells (Transgene biotech, Beijing, China). Each gene selected 3 positive clones for sequencing.

### Tissue-specific expression analysis

cDNA samples synthesized from total RNA from leaves, styles and pollen of the 37 peach varieties included in this study were used as templates to analyze tissue-specific expression of *PperS-RNases, PperSFBs* and *PperSLFLs*. Gene specific primers were designed and listed in Supplemental Table S2, and an actin gene was used as an internal control for constitutive expression with the following thermal cycling condition: a denaturation step at 94°C for 5 min followed by 30 cycles of 95°C for 30 s, 60°C for 30 s. 72°C for 30, and then 72°C for 10 mins.

### Construction of the phylogenetic tree of F-box, CUL1 and SSK1

64 CDSs of S-locus F-box genes from *Malus domestica, Pyrus pyrifolia, Pyrus bretschneideri, Prunus avium, Prunus dulcis, Prunus mume, Prunus salicina, Prunus armenica* were used to construct phylogenetic trees with the f-box genes specifically expressed in pollen cloned from 37 peach varieties in this study. The deduced amino acid sequences of Skp1-like proteins and cullin-like proteins from *Arabidopsis thaliana, Antirrhinum hispanicum, Prunus tenella, Petunia integrifolia, Pyrus bretchneideri, Malus domestica, Prunus avium, Prunus persica. Petunia integrifolia, Vitis vinifera, Nicotiana tabacum, Prunus tomentosa* and *Prunus mume* were aligned by CLUSTALW. Based on the alignment, phylogenetic trees were constructed by the neighbor-joining method (Saitou and Nei, 1987) using MEGA 6.0 program with the neighbor-joining method and the bootstrap test replicated 1000 times. The confidence values were shown on the branches.

### Yeast Two-hybrid (Y2H) Analysis

Yeast transformation and activity of β-galactosidase assays were performed following the manufacturer’s instructions (Clontech, CA, USA). The partial CDSs of *PperS-RNases* removed signal peptide and the full-length CDSs of *PperSSK1* and *PperCUL1* were cloned into pGBKT7 vector (Clontech), whereas the full-length CDSs of *PperSFBs, PperSLFLs, PperSSK1* and *PperRbx1* were cloned into pGADT7 vector (Clontech).

In order to find out the reason that PperSFB differentiates between self S-RNase and non-self S-RNases, Normal *PperSFB2* was divided into four parts: box, box-V1, V1-V2 and HVa-HVb and cloned them into pGADT7 vectors. Yeast two-hybrid assay was performed to observe the interactions between different portions of PperSFB2 and all PperS-RNases in this study.

For the Y2H assay, AH109 cells containing both AD and BD plasmids were grown on SD/-Leu/-Trp medium for 3 d at 30°C. Ten independent clones for each combination were streaked on SD/-adenine/-His/-Leu/-Trp medium and grown for 3-4d at 30 °C. Then use X− a-gal (TaKaRa Bio) to dye the clones to determine if there were interactions between the combinations. For quantitative measurements, β-galactosidase activity was determined using o-nitrophenyl-β-D-galactopyranoside (Sigma Aldrich) as a substrate according to Hao J. (2009) described.

### Bimolecular fluorescence complementation (BiFC) analysis

The pCambia1300 vector was used to construct BiFC vectors, which contained the N− or C-terminal of yellow fluorescence protein (YFP) fragments (YFPN and YFPC), respectively. The full-length CDSs of *SLFLs* without stop codon were cloned into pCambia1300-YFPN vectors, whereas the partial CDSs of S-RNases without stop codon removed signal peptide were cloned into pCambia1300-YFPC. All the construct vectors were transformed into Agrobacterium tumefaciens GV3101 and co-infiltrated into *Nicotiana Benthamiana* leaves. Fluorescence was observed in epidermal cell layers after 5 days by Olympus BX61 fluorescent microscope (Olympus FluoView FV1000).

The box, box-V1, V1-V2 and HVa-HVb frames without the stop codon were amplified and cloned into the pCambia1300-YFPN vectors. The recombinant plasmids containing the *box-YFPN, box-V1-YFPN, V1-V2-YFPN* or *HVa-HVb-YFPN* fusion gene and *PperS1-RNase, PperS2-RNase, PperS2m-RNase* or *PperS4-RNase* removed signal peptide fusion gene and the control plasmid with *YFPN* and *YFPC* were co-transformed into maize (*Zea mays* Linn.Sp.) protoplasts respectively according to Ren et al. (2011). GFP fluorescence was observed by Olympus BX61 fluorescent microscope (Olympus FluoView FV1000).The primers used were listed in (Supplemental Table 2).

### Purification of Peptide Tagged Recombinant Proteins

His-tagged proteins were purified as previously described (Meng et al., 2014). The mature peptide of the S-RNases without signal peptide were cloned into the pEASY-E1 vector (From TransGen Biotech Company) and transformed into the *E. coli* strain BL21 plysS (DE3) (From TransGen Biotech Company). The cells were inoculated into LB medium containing 100μg/ml ampicillin and incubated for about 3h at 37 °C in a shaker at 200 rpm. Once the cell suspension OD600 reached about 0.5, isopropyl β-D-1-thiogalactopyranoside (IPTG) was added into medium with final concentration of 0.2 mM to induce protein expression. The cell suspension was incubated for 10-12 h at 16°C in the shaker at 180rpm. His-tagged fusion proteins were purified using Ni-NTA His Binding resin (Novagen, USA) as previously described (according to the manufacturer’s instructions of Ni-NTA His Bind Resins, Novagen). The full-length coding sequences of pollen-expressed *PperSFB1m, PperSFB2m, PperSFB2, PperSFB4m PperSLFL1, PperSLFL2* and *PperSLFL3* were cloned into pMAL-c5x vector, which is designed to generate maltose-binding protein (MBP) fusion proteins. Similarly, the *PperCUL1, PperSSK1* and *PperRbx1* were cloned into pGEX4T-1 vector, which is designed to produce glutathione S-transferase (GST) fusion proteins. All the GST-fusion proteins and MBP-fusion proteins were purified using glutathione resin and maltose, as previously described (Yuan et al., 2014).

### *In Vitro* Ubiquitination Analysis of S-RNase

*In vitro* ubiquitination assays were performed as previously described (Yang et al., 2009; Yuan et al., 2014). The reaction mixture containing 50 mM Tris (pH 7.4), 10 mM MgCl_2_, 2 mM dithiothreitol (DTT), 5 mM HEPES, 2 mM adenosine triphosphate (ATP), 0.05% Triton X-100, 10 mM creatine phosphate, 1 unit of phosphokinase, 10 µg ubiquitin, 50 nM E1 (UBA6, *Petunia hybrida)*, 1 mM PMSF, 850 nM E2 (UBH6, *P. hybrida*), and aliquots of the recombinant proteins GST-PpSSK1, GST-CUL1, GST-Rbx1 and any MBP-SLFL at 30 °C for 2 h. Mixtures were immunoblotted using an anti-S-RNase antibody (From Beijing ComWin Biotech Company).

## Results

### Identification of *PperS-RNase* and *PperSFB* alleles in 37 Chinese peach varieties

For this study, we collected 37 Chinese peach varieties which represent local cultivars in 18 provinces/municipalities in China (Supplemental Table 1). These are thought to represent the evolutionary paths from the origin in central China (Tibet, Yunnan and Guizhou provinces) to the northwest of China (Shanxi province), then to the southwest of China and finally to the coastal and Xinjiang provinces (Cao et al., 2014) (Supplemental Fig. S1). Only four *S*-haplotypes *S*1, *S*2, *S*2m and *S*4, were detected from 36 peach varieties except Guang He Tao (Supplemental Fig. S2) (Supplemental Table 1), and they had previously been reported (Tao et al. 2007). The S genotype of 18 varieties, including Da Hong Pao, were S2S2 genotype and 9 varieties, including Hunchun Tao, were S1S2 genotype, while 3 varieties were of the S2S4 genotype (Feicheng Bai Li 10, Feicheng Bai Li 17 and Feicheng Hong Li 6) (Supplemental Table 1). The *S2-RNase* in 6 varieties was observed to contain a nucleotide substitution (G--A), which results in the conversion of the sixth conserved cysteine residue to a tyrosine in the *Prunus* C5 domain (Fig. 1A). This was named as *S2m-RNase. S2-RNase* and *S2m-RNase* were also identified from the original species Guang He Tao, which indicating that the mutation of *S2-RNase* had occurred before the formation of Chinese peach cultivars. Interestingly, we found that two *S-RNases* cloned from Guang He Tao lacked two introns (Fig. 1A). The 4 *S-RNases* were only expressed in pistil (Fig. 1D).

**Fig. 1.**
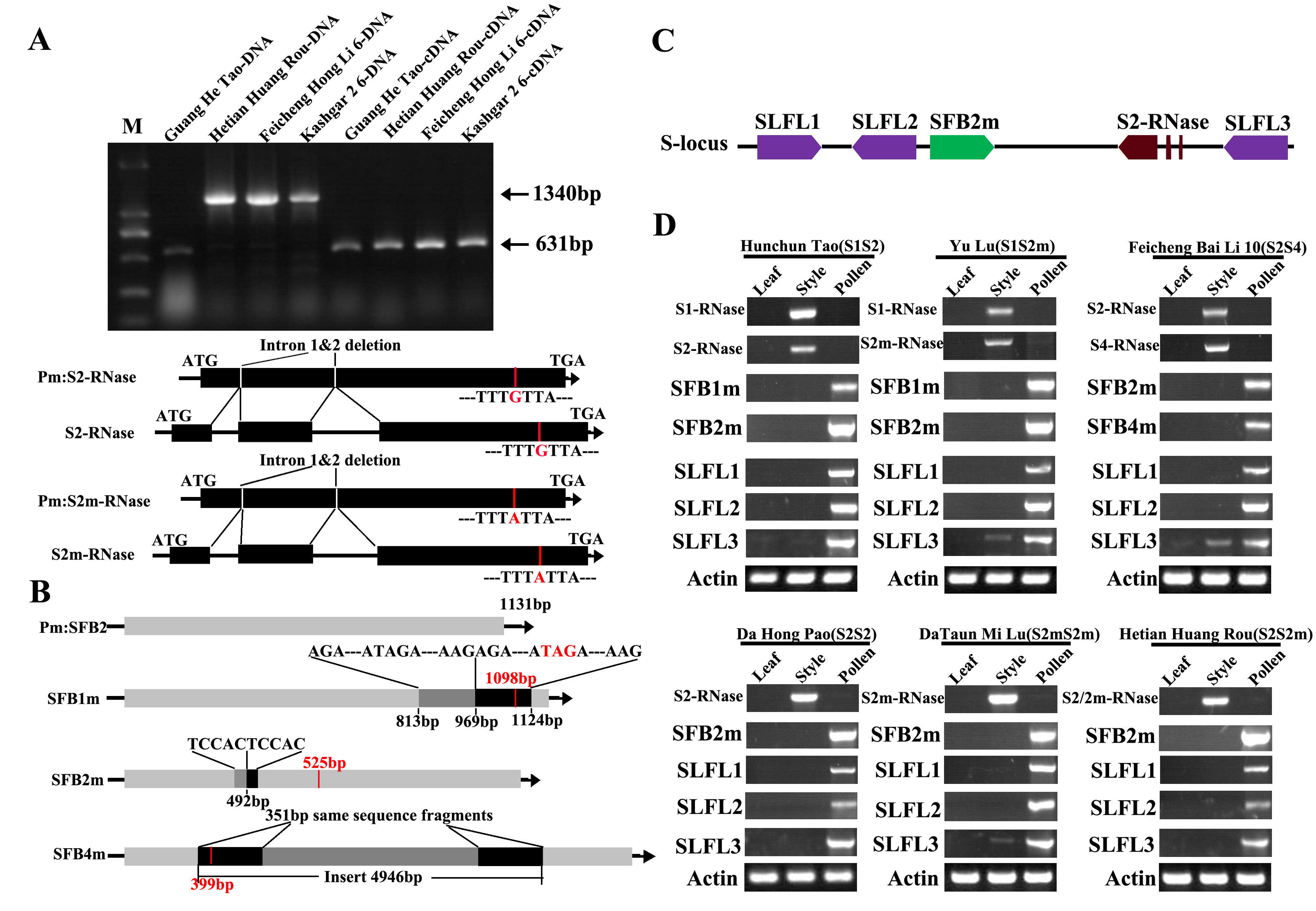
Characterization and expression patterns of Chinese peach *S* genes. (A) PCR analysis and schematic diagrams of the S2/2m-RNase in ‘Guang He Tao’’. The red line represents the mutation site of the S2-RNase. (B) Schematic diagram of Chinese peach SFBs. The black arrows indicate the transcriptional orientations of the genes. The red vertical bars indicate the stop codon, and the red numbers represent the length of the encoding frame. The black boxes in SFB1m and SFB2m represent the inserted fragment, and the gray boxes represent the same fragment of the gene as the inserted fragment. The black boxes represent the same fragments at both ends of the inserted fragment. (C) Schematic diagrams of the location of S2-RNase, SFB2m and SLFLs at the S-locus. The directions of the arrows represent the transcriptional orientations of S2-RNase, SFB2m and SLFLs. The middle parts in the red box represent the introns. (D) Tissue-specific expression analysis of *S1-RNase, S2-RNase, S2m-RNase* and *S4-RNase, SFB1m, SFB2m, SFB4m* and *SLFLs.* Total RNA from different organs was extracted and used as template for cDNA synthesis.

The mutated *PperSFBs* cloned from all varieties in this study were the same as previously reported mutations, but we found that the sequence of inserted 155 bp fragment in *PperSFB1m* was the same as the sequence of 155 bp fragment upstream of the insertion point, and the sequence of 5bp insertion in *PperSFB2m* was also the same as the 5 bases upstream of the insertion point. The sequences of 351 bp at both ends of the inserted 4949 bp fragment in *PperSFB4m* were also the same (Fig 1B). The mutant repeat sequences of *PperSFB4m* had previously been reported (Toshio et al., 2014). In addition, except the mutated *SFB2m*, a canonical *SFB2* gene was cloned from Guang He Tao, indicating that peach mutations occurred prior to evolution and the canonical *SFB2* was eliminated during the selection process. The expression of *SFBs* was pollen-specific (Fig. 1D).

### Cloning and expression analysis of *PperSLFL* and SCF complex (*PperCUL1*, *PperSSK1* and *PperRbx1* genes)

According to the sequences of *S-locus F-box-likes (SLFLs*) in peach genome (Genome Database for Rosaceae, http://www.rosaceae.org/), the primers were designed to clone the six *SLFL* genes. Finally, only three *SLFL* genes were cloned from pollen cDNA and their expression was pollen-specific (Fig. 1D). We named the three F-box genes as *PperSLFL1, PperSLFL2* and *PperSLFL3*, respectively. *PperSLFL1* located at about 47kb downstream of *PperS2-RNase* and the translation direction was opposite to that of *S2-RNase; PperSLFL2* located at about 26kb downstream of *S2-RNase* and *PperSLFL3* located at about 1.3kb upstream of *S2-RNase*, and the translation directions of the both F-box genes were the same as that of *S2-RNase.* The three *PperSLFL* genes did not have introns (Fig. 1C). The *PperSLFL1* and *PperSLFL2* showed pollen-specific expression, similar to the *PperSFB* genes. Despite the fact that *PperSLFL3* transcripts were detected in both leaves and styles, its expression was the most in pollen (Fig. 1D). The identity of the predicted amino acid sequences of PperSLFL1, PperSLFL2 and PperSLFL3 was 52.45%, while the alignment of the predicted amino acid sequences of PperSLFL1 with PperSLFL2 and PperSLFL3, respectively, the identity was 31.33% and 31.72%, respectively. The 3 *PperSLFL* genes showed low sequence identity with each other and with *PperSFB.* All the *PperSLFL* proteins contained the basic F-box domain and the FBA domain (Fig. 2). The phylogenetic tree analysis showed the three *PperSLFL* genes of peach clustered together with other *Prunus SLFL* genes, which was reported previously (Tao et al., 2008). The phylogenetic tree had two large lineages; the *Prunus SFB* genes did not cluster together with the pollen S genes of Pyrus and Malus and diverged into a separate lineage. Pyrus and Malus *SFBBs* and *Prunus SLFL* genes clustered together and diverged into two secondary lineages. *Prunus SLFL* genes were more closely related to Pyrus and Malus *SFBB* genes than *Prunus SFB* genes (Supplemental Fig. S3A).

**Fig. 2.**
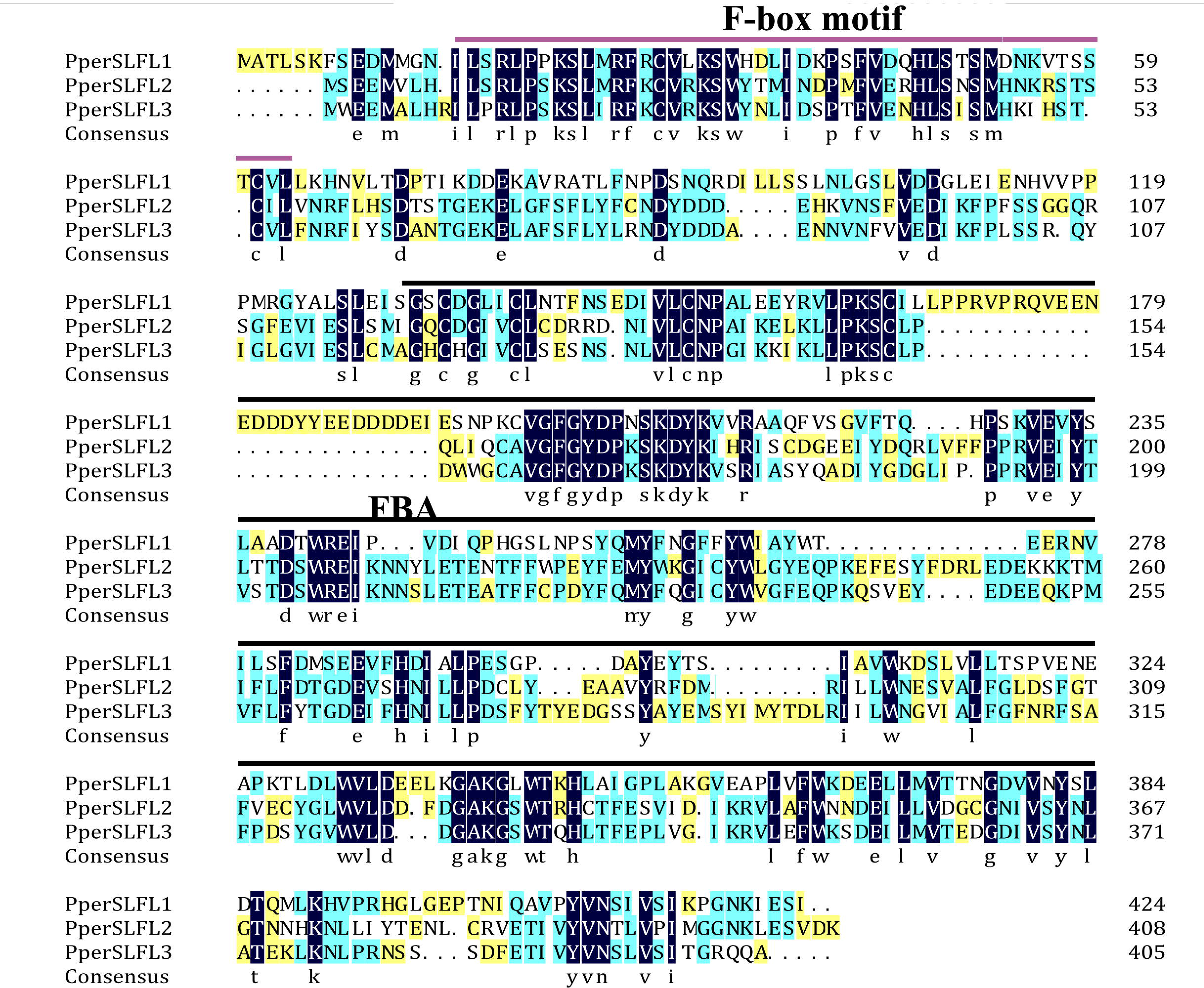
Aligment of the deduced amino acid sequences of PperSLFLl, PperSLFL2 and PperSLFL3. The three PperSLFLs sequences of Peach aligned using DNAMAN. The F-box domain is marked by purple line above, and the FBA domain is marked by black line above.

The SSK1 proteins with two conserved domains of apple, pear and petunia were used in a BLAST search for the proteins predicted in the peach predict protein database (http://www.rosaceae.org/). The candidate *SSK1* gene (*PperSSK1*) was cloned with specific primers (Supplemental Table 2). The full-length coding sequence (CDS) of *PperSSK1* was amplified from ‘Hunchun Tao’ (*S1S2*) pollen and was subsequently identified in the other 36 peach varieties. The canonical Skp1 comprises 150 to 200 amino acid residues and contains a Skp1-POZ domain at the N terminus and a Skpl domain at the C terminus. The deduced amino acid sequence of PperSSK1 comprised 177 residues and contained the Skp1-POZ and the Skp1 domain. In the phylogenetic tree, PperSSK1 clustered into a lineage with PtSSK1 and PavSSK1 (Supplemental Fig. S3B). In addition, the other two subunits of the SCF complex, *PperCUL1* and *PperRbx1*, were cloned with pollen cDNA of ‘Hunchun Tao’ (*S1S2*) as template. The candidate *PperCUL1* gene encoded the protein containing 744 amino acid residues and phylogenetic analysis showed that it clustered together with MdCULl and PavCULB which have been shown to be a component of the SCF complex (Tao et al., 2016; Yuan et al., 2014) (Supplemental Fig. S3C). PperRbx1 protein contained 117 amino acid residues, and had an H2 loop figure domain at the C-terminus which is necessary for ubiquitin ligase activity.

### The Interactions between S-RNases and S-locus F-boxes

Yeast two-hybrid analysis was performed to detect the interactions of PperSFBs with S-RNases. *PA1* (ppa011133m) (Aguiar et al., 2015), also a T2-RNase of *Prunus persica*, was cloned in this study. Because of no signal peptide, the full-length coding sequence (CDS) of *PA1* was cloned into pGBKT7 to detect the interactions of PA1 with S-locus F-boxes. The results showed that the mutated PperSFB and normal PperSFB2 interacted with all the S-RNases, and these interactions displayed no allelic specificity. There was no interaction between PA1 and PperSFB (Supplemental Fig. S4A). Furthermore, the (β-galactosidase reporter gene activity was detected and it showed that the intensity of interactions between these combinations was not high and the intensity of interaction between normal PperSFB2 and S2-RNase was slightly higher than that of other combinations (Supplemental Fig. S4B). Because of the insertion of the fragments, the proteins encoded by *PperSFB1m, PperSFB2m* and *PperSFB4m* genes were terminated prematurely and the domains at C-terminus was lost in varying degrees. In order to explore the effect of each part of the SFB on S-RNase, we divided the normal *PperSFB2* gene into four parts: box, box-V1, V1-V2 and HVa-HVb (Supplemental Fig. S4C). Using yeast two-hybrid and bimolecular fluorescence complementation (BiFC) analysis, we found that the box region did not interact with all of four S-RNases, whereas box-V1 and V1-V2 portions of PperSFB2 physically interacted with all four S-RNases, and the interactions displayed no allelic specificity. From the yeast coloring time stained by X-α-gal and fluorescence intensity, it could be seen that the interaction intensity between each combinations was not high. Interestingly, the HVa-HVb of PperSFB2 only interacted with S2-RNase but not other S-RNases, indicating a potential role in S-RNase-SFB specific recognition (Supplemental Fig. S5).

The expression of *PperSLFL1-3* had pollen specificity and contained F-box domains, which led to a problem that whether they might play roles in self-incompatibility of peach. First, we detected the interactions of PperSLFL1-3 with S-RNases. The Y2H and BiFC analysis were performed using various fusion expression vectors containing *S-RNases* or *PperSLFL1-3*, respectively. The Y2H assay showed that S-RNases interacted with PperSLFL1-3 with no allelic specificity, respectively (Fig. 3A). The β-galactosidase report gene activity suggested that the intensity of interaction between PperSLFL1 and the four S-RNases was slightly higher than other various combinations (Fig. 3E). PA1 did not interact with any PperSLFL proteins (Fig. 3A). To confirm that results, bimolecular fluorescence complementation (BiFC) experiment was performed in *Nicotiana tabacum* leaves. The BiFC results also indicated that PperSLFL1-3 interacted with the four S-RNases with no allelic specificity (Fig. 4).

**Fig. 3.**
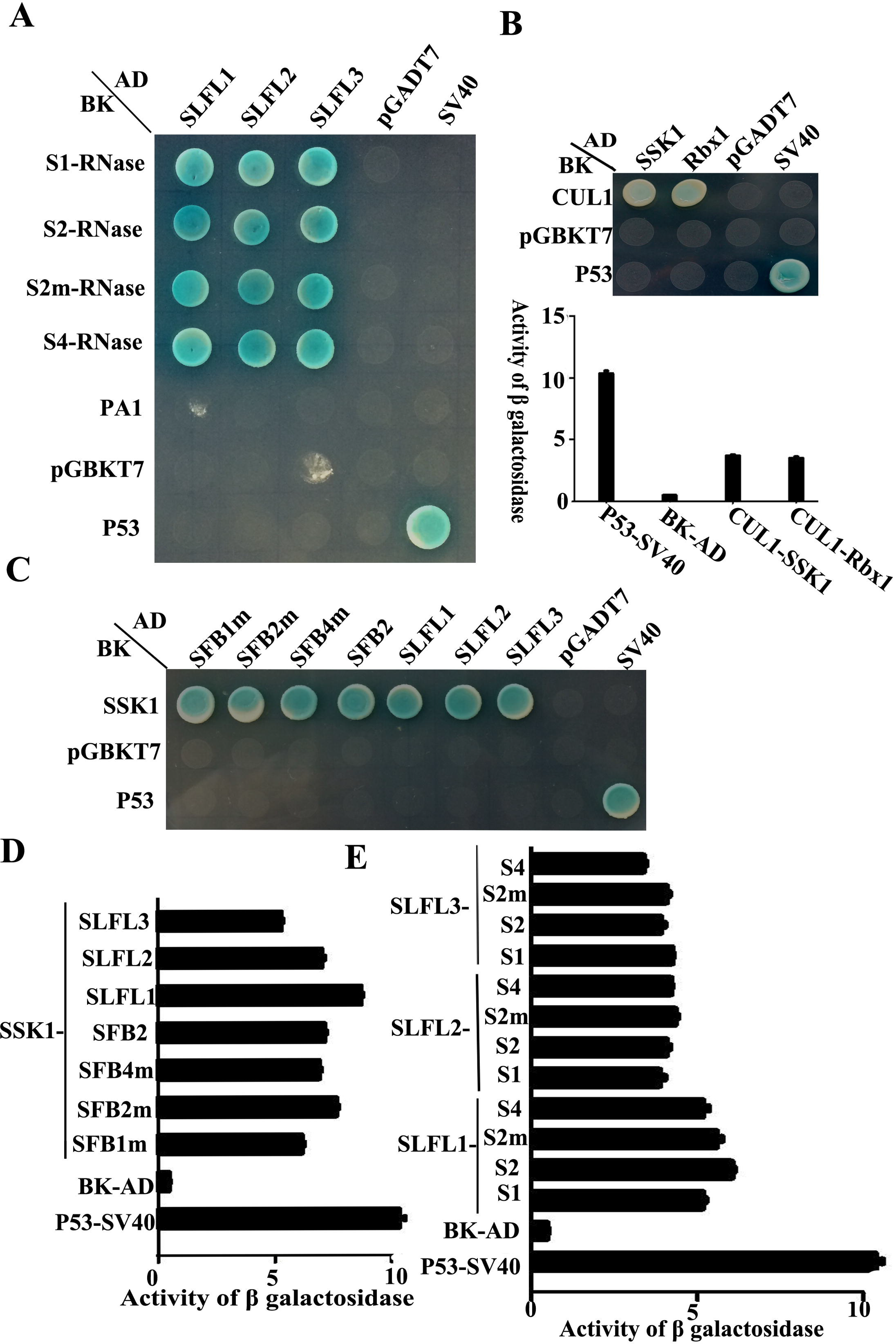
Yeast two-hybrid (Y2H) analysis to investigate interaction between S-RNases, PperSFBs, PperSLFLs, PperSSK, PperCUL1 and PperRbx1. (A) and (E) Y2H assays and the activity of β-galactosidase assay of the interaction between PperSLFLs and S-RNases, PA1 (ppa011133m, a T2-RNase in *Prunus persica.*) (B) Y2H assays and activity of β-galactosidase assay of interaction between PperSSK1, PperRbx1 and PperCUL1. (C) and (D)Y2H assays and the activity of β-galactosidase assay of interaction between PperSSK1 and PperSFBs and PperSLFLs. Empty vector was used as negative control; SV40 and p53 was used as positive. AD-activation domain; BD-DNA binding domain.

**Fig. 4.**
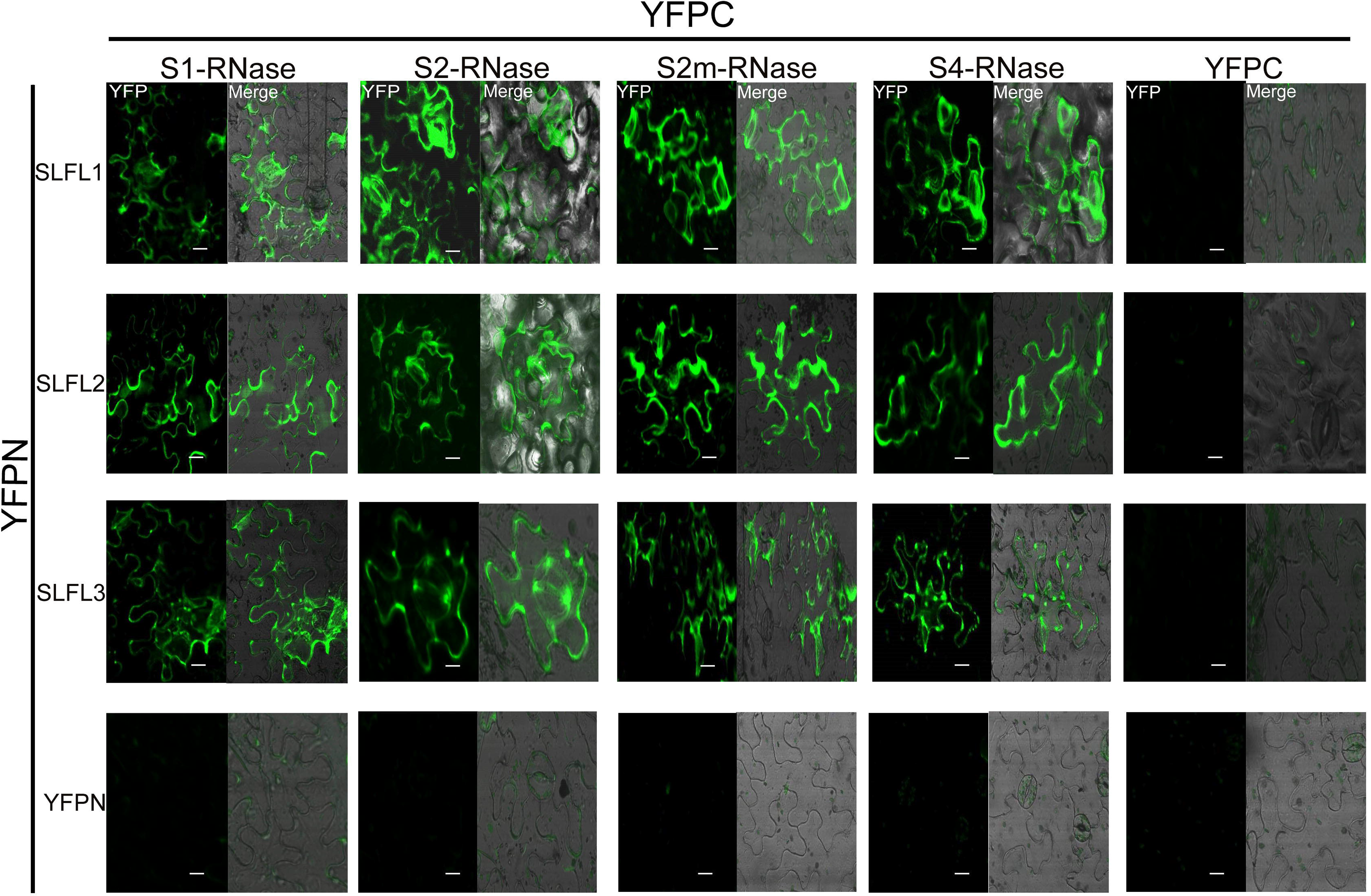
Bimolecular fluorescence complementation (BiFC) analysis of interactions between PperSLFLs with S-RNases. Construct pairs of PperSLFLs-YFPN, S-RNases-YFPC, YFPN and YFPC were transiently co-expressed in *Nicotiana tabacum* leaves. Fluorescence is indicated by the YFP signal. Merged images of YFP as well as bright field images are shown. PperSLFLs-YFPN and S-RNases-YFPC were co-injected with empty vector respectively as negative control. Scale bars = 10 μm.

### Interaction analysis of S-locus F-boxes, PperSSK1, PperCUL1 and PperRbx1

The yeast two-hybrid analysis was performed to investigate the interaction between PperSSK1 and PperSFBs/PperSLFLs, and the interaction between PperCUL1 and PperSSK1/PperRbx1. The results indicated that PperSSK1 interacted with PperSFB2, PperSFB1m, PperSFB2m and PperSFB4m (Fig. 3C), and the activity of β-galactosidase confirmed the intensity of the interaction was high (Fig. 3D). Both PperSSK1 and PperRbx1 interacted with PperCUL1, and the activity of β-galactosidase analysis quantitatively demonstrated the interaction between them (Fig. 3B).

We also examined the interactions between PperSSK1 and three other pollen-expressed F-box proteins, PperSLFL1–3, to explore the hypothesis that PperSLFL proteins might play a role in self-incompatibility of peach. The yeast two-hybrid analysis was conducted using the full-length CDSs of these 3 S-locus F-box protein genes. All the three F-box proteins showed interaction with PperSSK1 (Fig. 3C). The activity of β-galactosidase report gene quantitatively demonstrated the interaction between them, and the interaction between PperSSK1 and PperSLFL1 and PperSLFL2 was stronger than that of between PperSSK1 and PperSLFL3 (Fig.3D).

### A SCF complex containing PperSLFL1-3 ubiquitinates S-RNases in vitro

In order to test whether *S*-RNases could be ubiquitinated by SCF^SLFL^ in vitro, commercial His-UBA6 was used as the ubiquitin-activating (E1) and His-UBH6 was used as ubiquitin-conjugating enzyme (E2). Purified MBP-PperSLFL1-3, GST-PperSSK1, GST-PperRbx1 and GST-PperCUL1 were used as E3. Anti-MBP antibody was used to detect MBP-PperSFB proteins and MBP-PperSLFL proteins by Western blot analysis, anti-S-RNase antibody was used to detect S-RNase proteins and ubiquitinated *S*-RNases and anti-GST antibody was used to detect GST-tagged proteins. Purified His-S-RNase proteins, MBP-SFB proteins and MBP-SLFL proteins were detected a single band (Fig. 5). Distinct immunoreactive bands with higher molecular masses (between 34 KDa and 130 KDa) were detected above the predicted His-S-RNase (26 KDa) bands, and no band was detected in the negative control reactions without His-S-RNase proteins (Fig. 6). The results indicated that PperSLFL proteins could ubiquitinate *S*-RNase of peach.

**Fig. 5.**
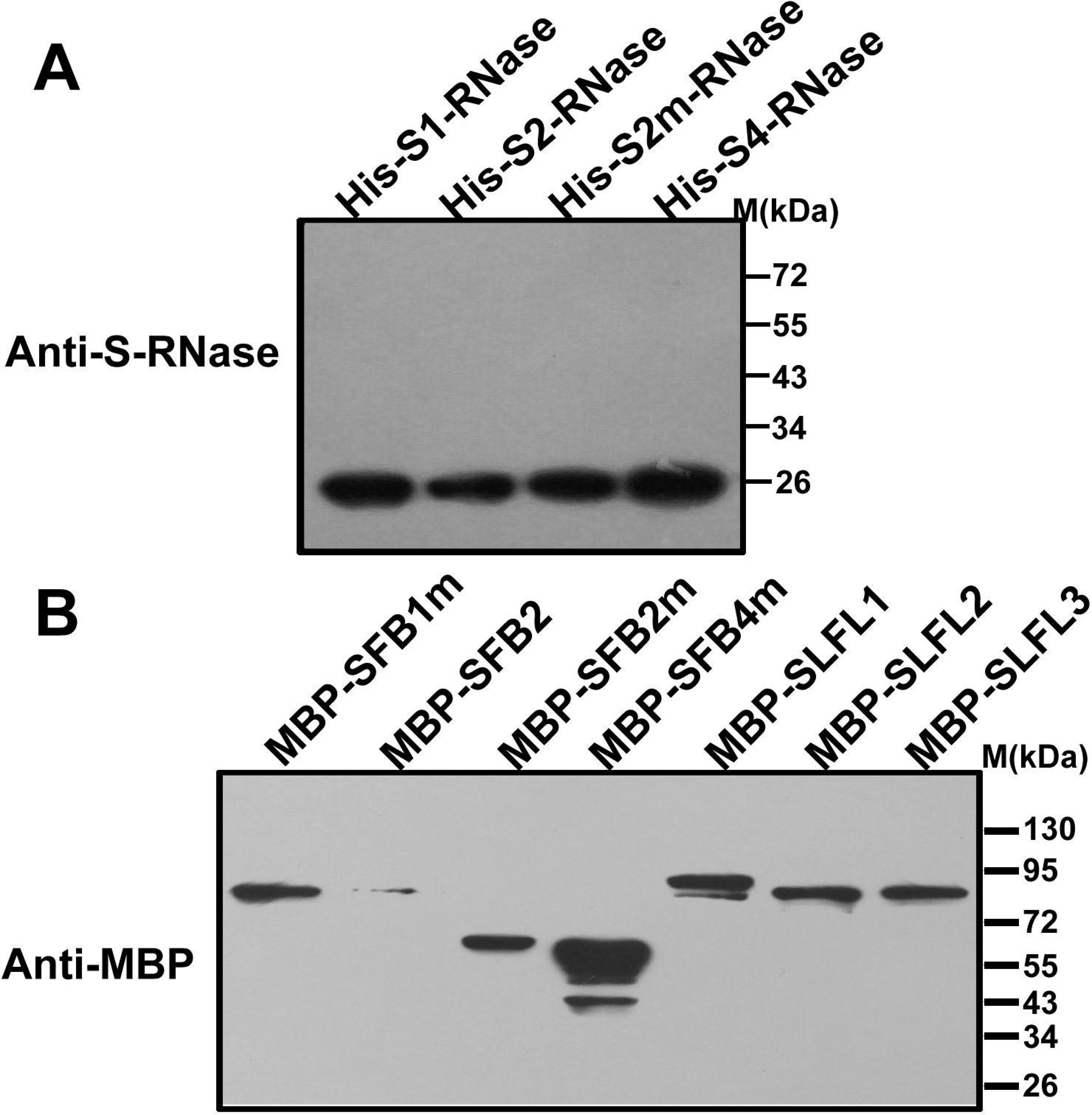
Immunoblot detection of S-RNases and F-box proteins. *E.coli* expressing S-RNases were detected by a polyclonal antibody. The polyclonal antibody against the recombinant S2-RNase was raised in rabbit and the antibody detected not only S2-RNase but also other allelic S-RNases without allelic specificity. *E.coli* expressing F-box proteins were detected by commercial mouse monoclonal antibody. 10 μg of total proteins were loaded in each lane.

**Fig. 6.**
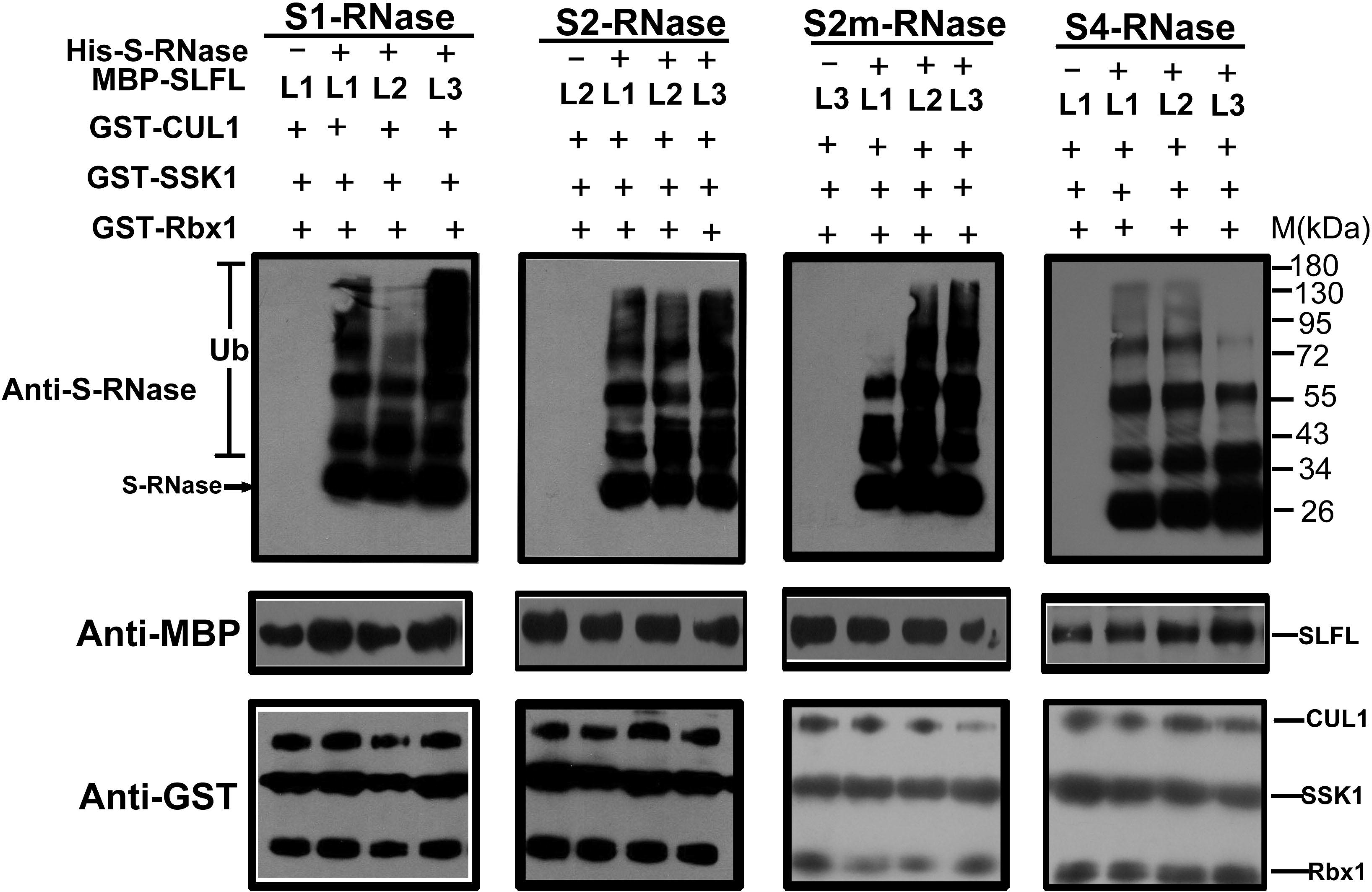
*In vitro* detection of ubiquitinated S-RNases. Each of the lanes was loaded with 10 μg protein. The ubiquitination of different PperS-RNases was analyzed with the presence of PperSSK1, PperCUL1, PperSLFL1-3 and Ub. 10 μg of PperSLFL1-3 proteins was added to each lane, respectively. The lanes without His-tagged S-RNase were used as negative control.

## Discussion

In this study, we selected 37 Chinese peach varieties from 18 areas of China, including the ancestral species (Guang He Tao), wild species (Qing Si, Huo Lian Jin Dan, Qing Mao Zi Bai Hua, Bai Nian He, Zhang Bai 5 and Long 1-2-4) and some common local varieties. After identifying the S-genotypes of all the peach varieties in this study (Supplemental Table 1), four previously described S haplotypes were identified S1, S2, S2m and S4. S2 was the most frequent S-haplotype in the tested Chinese peach varieties (occurred in 33 varieties), followed by S1 (in 10 varieties), S2m (in 7 varieties) and S4 (only in 3 varieties). All four S haplotypes, S1 S2, S2m, S4, found in this study had the same mutant versions as that reported previously (Tao et al., 2007; 2010). The mutated *S2m-RNase* and *SFB2m* genes, canonical *SFB2* gene and *S2-RNase* existed in Guang He Tao indicating that these mutations occurred before the formation of Chinese peach cultivars. In the mountains of Tibet, peach might be propagated by seeds generally obtained from genotypes with high productivity. Because of the special natural environment, self-compatibility generally led to more reliable fruit set, which made the probability of survival in natural selection increased, but the genetic diversity in the S-locus was declined. We proposed that under selection pressure for SC, pollen part mutants might preferentially be selected compared to pistil part mutants because there were many pollen grains and the pollen genotype in a large extent affected the SI phenotype in GSI system. That is why all the peach S haplotypes characterized in this study are pollen part mutant S haplotypes and the S2-allele accumulates the most.

After analyzing the sequences of mutant pollen *S* genes, we found that the 155bp fragment inserted in *PperSFB1m* was duplicated from the 155bp region upstream of the insertion point, and the 5 bp fragment inserted in *PperSFB2m* was the same with the 5 bp upstream of the insertion point. The sequence of 351 bp at both ends of the inserted 4949bp fragment in *PperSFB4m* was also the same that had been reported (Fig. 1B) (Tao et al., 2010). That kind of mutant might be due to an error in homologous recombination, or a retro-transposition. As we know, gene duplication is a ubiquitous biological phenomenon, an important driving force for the diversification of genomic and genetic systems, and plays a very important role in the evolution of biological processes. This repetition in peach might have a significant for the study of its evolution.

In the S-RNase-based GSI system, non-self S-RNases in pollen tube are detoxified. It is hypothesized that detoxification of non-self S-RNases in pollen is mediated by the SCF complex. S-RNases are degraded by SCF^SLF/SFBB^ mediating in other plant species with the S-RNase-based GSI system (Tao and Iezzoni, 2010; Iwano and Takayama, 2012; Yuan et al., 2014). In this study, three pollen-expressed F-box genes (Fig. 1D and Fig. 2), named as *PperSLFL1-3*, located at the S locus of peach (Fig 1C). Phylogenetic and evolutionary analysis indicated that PperSLFL1-3 clustered with SLFLs of other *Prunus* species on the same evolutionary branch, and the evolution relationship between SLFLs and SFBBs of apple and pear was closer than the evolution relationship between SLFLs and SFBs (Supplemental Fig. S3), which was the same as Tao did (Tao et al., 2008). We speculated that the *Prunus* SI recognition mechanism might have some differences compared to the mechanism in the Maloideae. Researchers speculated that *Prunus* self-incompatibility mechanism was ‘self recognition by a single factor’ system (Sonneveld et al., 2005). In the ‘self recognition by a single factor’ system, the cytotoxic effect of non-self S-RNases is thought to be inactivated by an unidentified ‘general inhibitor’ (GI) (Sonneveld et al., 2005). The single factor we speculated was SFB, because the peach became self-compatibility from self-incompatibility due to SFB mutant. SFB2 had a role in self/non-self recognition, the variable region interacted with self/non-self S-RNase, and the hypervariable region interacted with self-S-RNase (Supplemental Fig. S5). We suspected that SFB specifically recognizes self-S-RNase to leave self-S-RNase active, leading to the arrest of self pollen tube growth.

The F-box protein would be the good candidate for the GI. In sweet cherry, S-RNases were recognized by PavSLFL2 (Matsumoto, 2016). In this study, PperSLFL1-3 interacted with all the S-RNases with no allelic specificity (Fig. 3A), and phylogenic analysis demonstrated that SLFLs including PperSLFL1-3 were classified into the same clade as SFBB of the Maloideae. Since it is plausible that SI or the S locus of Prunus and the Maloideae share the same origin (Igic and Kohn, 2001; Steinbachs and Holsinger, 2002), we suspect that *SLFLs* are homologs of *SFBB.* During the evolution of SI in Prunus, SLFLs may lose their function in S haplotype-specific interaction, and may recruit SFB for S haplotype-specific interaction. All together, these results suggested that PperSLFL1-3 have appropriate characteristics to be the GI.

Matsumoto (2012, 2016) showed that SFB and PavSLFLs interact with a Skp1-like1 homolog that is proposed to be a component of the SCF complex involved in the polyubiquitination of proteins targeted for degradation. By Y2H experiment, we have known that PperSLFL1-3 could interact with PperSSK1 and participate in the formation of SCF complex. It is possible that the PperSLFL1-3 would participate in the degradation of S-RNase proteins in a process as with that SLF/SFBB proteins are involved in the degradation of non-self S-RNase proteins (Kubo et al., 2010; Kakui et al., 2011; Williams et al., 2014; Kubo et al., 2015). In vitro ubiquitination analysis, we found that PperSLFL1-3 could make all the PperS-RNase proteins in this study tag the polyubiquitin chain (Fig. 6). According to the result, we suspect the SLFL proteins act as GIs to target all S-RNases with no allelic specificity in pollen.

In conclution, our results suggested that PperSLFL1-3 were a subunit of SCF complexes, recognized all S-RNases taken up into pollen tube and mediated polyubiquitination of S-RNases. Because loss-of-function of SFB results in pollen-part SC of peach unlike that in Japanese pear, the role of SLFL genes in the SI has been the focus of attention. The Y2H assay and activity of β-galactosidase assay showed that there was a strong interaction between PperSLFL1-3 and PperSSK1 and PperS-RNases. We thought that when S-RNases were taken up into the pollen tube, SFB would recognize self S-RNase and protect it by some kind of motification, and SLFL proteins could not recognize and target it. Cytotoxic effect of self S-RNase arrests pollen tube growing. When the SFB mutated, the ‘protection’ on self S-RNase disappeared, SLFL proteins target S-RNase and tag polyubiquitin chain on it, S-RNase could be degraded, and the pollen tube continue to grow to complete fertilization. This model is needed to be carfully tested, and further studies are needed to clarify the mechanism of self-incompatibility in *Prunus.*

## Supplemental data

**Supplemental Fig. S1 The distribution of 37 peach varieties in China** The arrows in the figure represent the evolutionary direction of peach in China. The red shade represents the origin of Chinese peach, and the gray shades represent the secondary center of origin of Chinese peach, and the purple circles represent various Chinese peach population.

**Supplemental Fig. S2 Identification for the S genotypes of 36 peach varieties except Guang He Tao** The PCR were performed with DNA extracted from leaves as template and primers Pru-C2/Pru-C4R for S_1_ and S_2_, and S_4_ specific primers.

Supplemental Fig. S3. **Phylogenetic trees of CDSs of S locus F-box and deduced amino acid sequences of Skp1-like proteins and cullin-like proteins**. (A) A neighbor-joining tree was constructed from 64 S locus F-box genes from *apple(Malus domestica;* MdSFBBs), *pear(Pyrus × bretchneideri;* PbSFBBs. *Pyrus pyrifolia*; PpSFBBs), sweet cherry(Prunus *avium*; PavSFBs PavSLFLs), almond(Prunus *dulcis*; PdSFBs and PdSLFLs), plum(Prunus *mume*; PmSFBs and PmSLFLs, *Prunus salicina*; PsSFBs), apricot(Prunus *armeniaca*; ParSFBs), sour cherry(Prunus *cerasus;* PcSFB26), peach(Prunus *persica;* PperSFBs and PperSLFLs) and *Prunus speciosa* (PspSFB1). (B) Skp1-like proteins and cullin-like proteins were used to construct NJ trees. The deduced amino acid sequences of Skp1-like proteins were from *Arabidopsis* thaliana(AtSKPs), *Antirrhinum* hispanicum(AhSSK1), *Prunus* tenella(PtSSK1), *Petunia integrifolia(PiSKP1,PiSKP3), Pyrus *×* bretchneideri(PbSKP1), Malus* domestica(MdSSK1-2), *Prunus* avium(PavSSK1) and *Prunus* persica(PperSSK1). The deduced amino acid sequences of cullin-like proteins were from *Arabidopsis* thaliana(AtCUL1-3), *Prunus* avium(PavCUL1A, PavCUL1B), *Petunia integrifolia* (PiCUL1C), *Vitis vinifera(VvCUL1-1,VvCUL1-2), Nicotiana* tabacum(NtCUL1-1) *Prunus* tomentosa(PtCUL1), *Prunus* mume(PmCUL), *Pyrusn × bretchneideri(PbCUL1*, PbCUL1-1), *Malus* domestic(MdCUL1-1,MdCUL1-2) and *Prunus persica* (PperCUL1). NJ trees were generated with 1000 bootstrap replicates.

**Supplemental Fig. S4 Yeast two-hybrid assay and for the interactions between PperSFBs and S-RNases** (A) Yeast two-hybrid assay for the interactions between PperSFBs and S-RNases. (B) The activity of β-galactosidase assay for the interactions between PperSFBs and S-RNases. Each of the combinations was selected 10 yeast plaques and then divided into 3 portions. Each portion was cultured and the activity of β-galactosidase was measured separately. (C) Constructs of AD::box, AD::box-V1, AD::V1-V2 and AD::HVa-HVb.

**Supplemental Fig. S5 Yeast two-hybrid assay and BiFC assay for the interaction between S-RNases and portions of PperSFB2** (A) Yeast two-hybrid assay for the interaction between S-RNases and portions of PperSFB2. Various combinations of AD and BD fusions are tested for their growth on SD/-Leu/-Trp/-His/-Ade media. (B) BiFC assay for the interaction between S-RNases and portions of PperSFB2. Fluorescence is indicated by the YFP signal. Merged images of YFP as well as bright field images are shown. Scale bars = 10 μm.

**Supplemental Table 1 Thirty seven wild and local genotypes of Chinese peach**

## Acknowledgements

We thank Professor Lirong Wang from the Zhengzhou Institute of Pomology for providing the peach material. This work was supported by the National Natural Science Foundation of China (31171941 and 31372035). We thank PlantScribe (www.plantscribe.com) for editing this manuscript.

## Contributions

T. L. and D. M. designed the study; Q. C., D. M. and Z. G. performed the experiment. W. L., X. D. Q. Y. and Y. L. contributed reagents/materials. T. L. and D. M. wrote the paper. All authors read and approved the final manuscript.

